# Adaptive suppression of threat-history stimuli

**DOI:** 10.1101/2025.06.09.658555

**Authors:** Jingqing Nian, Yu Zhang, Yu Luo

## Abstract

Previous studies have found evidence of adaptive suppression mechanisms for physically salient stimuli. However, it remains unclear whether a similar mechanism exists for threat-history stimuli. This study used a threat conditioning task to generate stimuli with and without a history of threat. In the subsequent visual search task, the spatial probability of distractors was manipulated to examine the influence of threat-history stimuli on distractor suppression. The results showed that distractors appearing at high-probability locations were effectively suppressed, and suppression was stronger for threat-history distractors than for no-threat-history distractors. These findings suggest that threat-history distractors are more effectively suppressed when they appear at a predictable location through an adaptive attentional suppression mechanism.

**Data availability:** All data supporting the findings are openly available via the Science Data Bank (https://www.scidb.cn/anonymous/VlpqNm55).

## 1 Introduction

In our daily life, the ability to detect and adapt to statistical regularities is fundamental to guiding attention and behavior. This form of implicit learning, known as statistical learning, allows individuals to predict events and allocate cognitive resources efficiently, even without explicit awareness(Fiser & Aslin, 2001; Turk-Browne et al., 2005). Recent research has demonstrated that statistical learning occurs across visual, auditory, and spatial domains and operates automatically in response to environmental structures(Fiser & Lengyel, 2022; Theeuwes et al., 2022).

Importantly, statistical learning plays a core role in shaping attentional processes(Theeuwes et al., 2022). While salient distractors typically capture attention (Lin et al., 2024; Theeuwes, 2010, 2025), a growing body of research has shown that the attentional capture of physically salient distractors (e.g., color singletons) can be reduced when these distractors frequently appear in a specific location within the visual field (Wang & Theeuwes, 2018a, 2018b, 2018c). Moreover, this suppression is saliency-dependent, with highly salient distractors more strongly suppressed at frequent locations(Failing & Theeuwes, 2020; Gong & Theeuwes, 2021). However, similar findings have not been consistently replicated in studies involving reward-related distractors(Kim & Anderson, 2021; Le Pelley et al., 2022), raising the question of whether threat-related stimuli follow a pattern more akin to reward or to physical salience. Previous studies suggest that threatening stimuli can capture attention more effectively than neutral stimuli, even when they are task-irrelevant(Anderson & Britton, 2020; Burra et al., 2019; Schmidt et al., 2015; Vuilleumier, 2005; Watson et al., 2019; Zsidó et al., 2023). This raises the question of whether motivational salient distractors associated with threat are also subject to adaptive spatial suppression, as is observed with physically salient stimuli.

Recent studies have begun to explore this issue by examining how motivational salience, particularly threat-related associations, interacts with distractor suppression(Theeuwes et al., 2025; Theeuwes & van Moorselaar, 2025). For example, Theeuwes et al. (2025) employed a two-phase design. In the initial pre-conditioning phase, participants developed learned suppression toward neutral distractors. In the subsequent post-conditioning phase, threat-associated distractors were presented at the same locations. The findings revealed that suppression remained stable across both phases, with no evidence that threatening distractors elicited stronger suppression at previously suppressed locations. One possible explanation is that suppression was already well-established during the pre-conditioning phase, which makes it unclear whether the observed effects in the post-conditioning phase reflected new learning or simply a continuation of earlier suppression. Notably, electric shocks were still administered on some trials during the post-conditioning phase to maintain the threat association. Although these trials were excluded from the analysis, the sustained emotional activation may nevertheless have disrupted the updating of suppression mechanisms(Pessoa, 2009; Anderson & Britton, 2020), thereby weakening the potential interaction between threat and location probability.

To address these potential limitations, the present study introduced two key modifications to the experimental design. First, the preconditioning phase was removed to allow direct examination of whether distractors with a history of threat affect learned distraction suppression, thereby avoiding confounds from prior suppression experience. Second, a classic fear conditioning paradigm was used to establish threat and non-threat stimuli. Then, an extinction phase was conducted to reduce emotional carryover effects. Before the visual search task, the shock electrodes were removed, and the participants were explicitly informed that no further shocks would be administered. This manipulation minimized the potential interference of emotional arousal on learning suppression. These adjustments provided a more robust test of adaptive suppression of threat-related influences. Specifically, we used a threat conditioning task to generate stimuli with and without a history of threat. In the subsequent visual search task, the spatial probability of distractors was manipulated to investigated whether the suppression of threat-history distractor is enhanced when it appear at high-probability locations during a visual search task.

## 2 Method

### 2.1 Participants

Participants were recruited from the subject pool of Guizhou Normal University. The study was approved by the Committee on Human Research Protection of the School of Psychology, Guizhou Normal University (IRB approval number: GZNUPSY.NO2022E [007]). All participants provided informed consent following the Declaration of Helsinki.

We used G*Power 3.1 to determine the required sample size to achieve 80% statistical power and an alpha level of 0.05. The minimum required sample size was 29 participants, based on an expected effect size of *d* = 0.54. This effect size was estimated from prior studies with similar designs (e.g., Kim & Anderson, 2021; Le Pelley et al., 2022). To account for potential exclusions, 33 participants were recruited. After excluding three individuals from the final analysis (see below for details), the final sample included 30 participants (17 females, aged 19–27 years), all of whom received monetary compensation for participating.

### 2.2 Apparatus

Screen luminance and color were measured and calibrated using the PsyCalibrator toolkit and SpyderX (Lin et al., 2023). The calibrated parameters were then applied during the stimulus presentation phase of the experiment. A Lenovo computer running MATLAB (version 2020b) and Psychtoolbox 3.0 generated the stimuli on a 21.5-inch LCD screen with a resolution of 1920×1080 pixels and a refresh rate of 60 Hz. All stimuli were presented on a black background (RGB: 5, 5, 5) from a distance of 72 cm.

### 2.3 Task, Stimuli, and Procedure

#### Shock work-up task

A multi-channel electrical stimulator (type: SXC-4A, Sanxia Technique Inc., Beijing, China) was attached to the left wrist of each participant and delivered stimulation through a pair of Ag/AgCl surface electrodes. The shock intensity was calibrated individually for each participant to reach a level rated as highly annoying yet not painful. The level was defined as a score of 8 on a subjective discomfort scale ranging from 0 (no feeling) to 10 (extremely painful).

#### Threat conditioning task

As visualized in Figure 1A, consistent with previous research(Bach et al., 2023), the threat conditioning task consisted of three phases: habituation, acquisition, and extinction. In each trial, one of two colors—green (RGB: 0, 73, 0) or cyan (RGB: 0, 71, 71)— was randomly assigned as the CS+ (paired with electric shock) or CS− (not paired with shock), with color-condition assignments counterbalanced across participants. Throughout all phases, participants were instructed to passively observe the stimuli without making any responses. The habituation phase included 4 trials: 2 CS+ trials (no shock) and 2 CS− trials. Each trial began with a 500 ms white fixation point, followed by an 8000 ms CS presentation. The acquisition phase consisted of 16 trials: 8 CS+ (6 with electrical stimulation, 2 without) and 8 CS−. During CS+ trials paired with shock, participants received an electric shock at the same time as the CS+ stimulus was presented. The trial structure was identical to the habituation phase, with inter-trial intervals (ITIs) randomly varying from 9000 to 15000ms. Electrical shocks in CS+ trials were administered at the intensity determined during the shock work-up task. No shocks were administered in the remaining CS+ or any CS− trials. The extinction phase included 40 trials: 20 CS+ and 20 CS−, all without electrical stimulation. The trial structure and ITIs remained consistent with the acquisition phase. The CS+ and CS− trials were presented in a randomized order throughout the entire task. After the acquisition phase, we removed the electrodes immediately and informed the participants that they would not receive any more shocks in the subsequent tasks.

**Figure 1.**
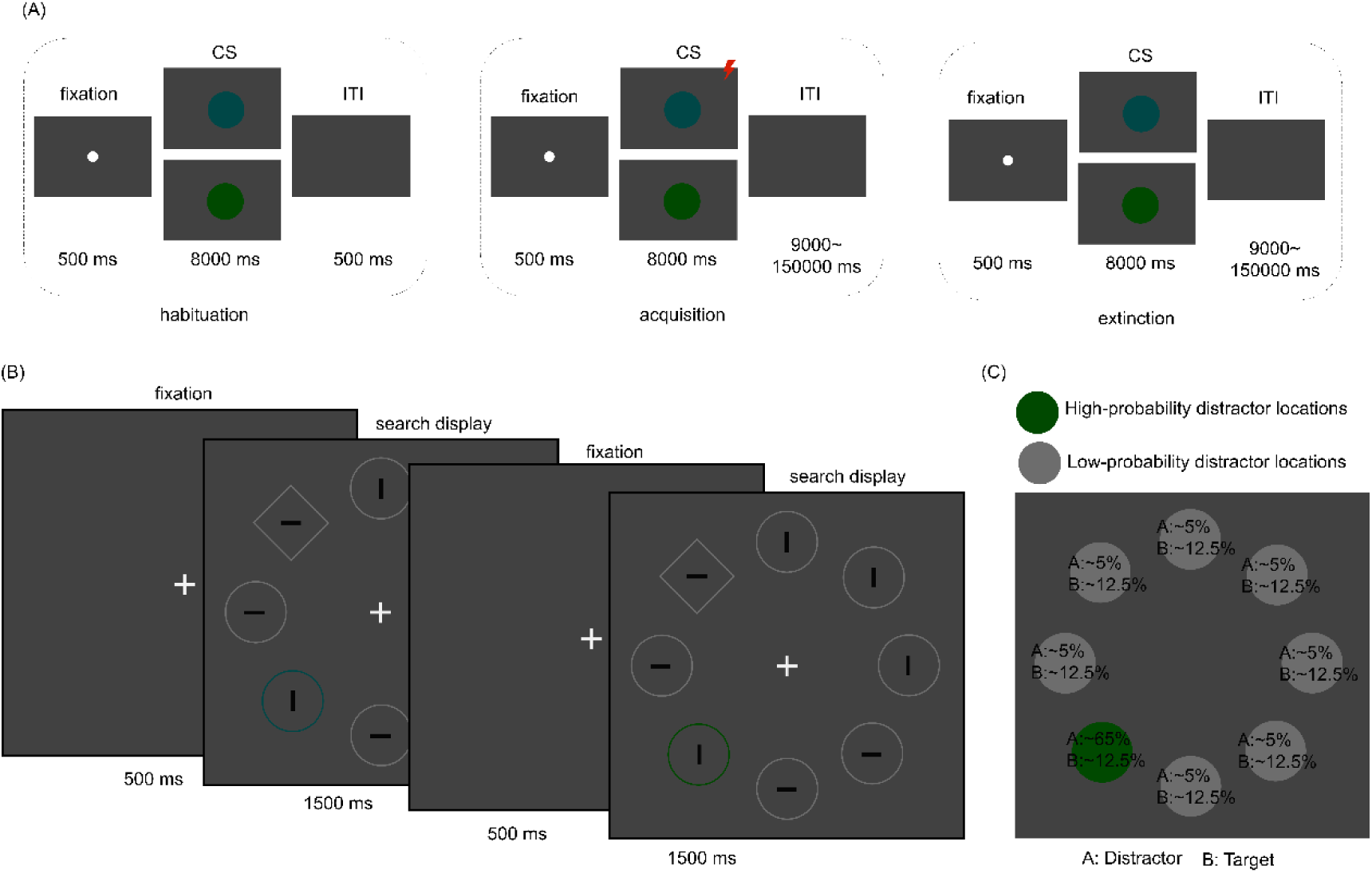
Experimental Paradigm. *Note.* (A) Example of the threat conditioning task. The task consisted of three distinct phases. *Habituation Phase*: Participants are exposed to the CS+ and CS− without any shocks to establish a neutral baseline. *Acquisition Phase*: The CS+ is followed by the shock on a majority of trials (i.e., 75%), while the CS− remains unpaired. *Extinction Phase*: Both the CS+ and CS− are presented again, but no shocks are delivered. This phase examines how quickly the learned fear response diminishes. (B)Example trial sequence of the visual search task. Each trial began with a fixation display, followed by a search display in which participants indicated the orientation (horizontal or vertical) of the line inside the target diamond. In two-thirds of the trials, one non-target shape was replaced by a singleton color distractor (either green or cyan). (C) Schematic representation of the spatial regularities of distractor presentation. Percentages at each location indicate the probability of the distractor or target appearing in that location throughout the task. See the online article for the color version of this figure.

#### Visual search task

As visualized in Figure 1B, each trial began with a central fixation cross presented on a black background. After 500 ms, a search display appeared and remained on screen until response or a 1500 ms timeout. The display contained eight equally spaced shapes (one diamond and seven circles), arranged on an imaginary circle (radius = 4.5°) centered on fixation. Each shape contained a gray line segment (RGB: 70, 70, 70) oriented either horizontally or vertically, randomly assigned. Participants reported the orientation of the line inside the target diamond by pressing either the s or k key, with key assignments counterbalanced across participants. They were instructed to respond as quickly and accurately as possible. In two-thirds of the trials, one non-target shape was replaced by a singleton color distractor (either green or cyan). The threat relevance of each color singleton (i.e., whether it had a threat history or no-threat history) was determined during the prior threat conditioning task. Critically, As visualized in Figure 1C, although the target appeared equally often in all locations, each color singleton distractor was more frequently presented at one specific location, defined as the high-probability distractor location (∼65%). The high-probability location was randomly assigned for each participant. Before beginning the formal task, participants completed 20 practice trials and were required to achieve at least 85% accuracy to proceed. The experimental session consisted of 16 blocks, each containing 56 trials: 20 trials with threat-history distractors, 20 with no-threat-history distractors, and 16 distractor-absent trials.

#### Post-task Awareness Assessment

After completing the visual search task, participants answered a series of questions assessing their awareness of the spatial distribution of distractors. They were first asked whether they had noticed any spatial imbalance in distractor occurrence. Then, they were required to indicate the location with the highest distractor probability for both the threat-history and no-threat-history distractors.

### 2.4 Data recording and Statistics

Participants’ responses were recorded during the visual search task, and the behavioral data were analyzed in R using RStudio. Accuracy (ACC) was examined using generalized linear mixed-effects models (GLMMs) and response times (RT) were examined using linear mixed-effects models (LMMs). Both were implemented with the *lme4* and *lmerTest* packages (Kuznetsova et al., 2017).

Prior to RT analysis, RT data were preprocessed according to the following steps: (1) The first two trials of each block were excluded, (2) Trials with incorrect responses were removed, (3) Trials with RTs shorter than 200 ms were excluded, (4) Trials were excluded if a participant’s reaction time in a given condition deviated by more than ±2.5 standard deviations from their mean response time in that condition, and (5) Participants with fewer than 70% of total trials remaining after preprocessing were excluded from further analysis. After the above preprocessing, the proportion of trials removed ranged from 7.48% to 26.79%, with a mean of 13.02% ± 0.89%.

In distractor-absent trials, both ACC and RT were modeled with distractor location probability (LocP) as a fixed effect and subject as a random intercept (model: ACC/RT ∼ LocP + (1 | Subject)). In distractor-present trials, both location probability and distractor category (CS) were included as fixed effects with subject as a random intercept (model: ACC/RT ∼ CS * LocP + (1 | subject)). Post-hoc comparisons were conducted using the *emmeans* package with Tukey adjustments for multiple comparisons.

To further assess how the development of the learned suppression effect differed between threat-history and no-threat-history distractors, we calculated the proportion of the interference reduction (*P*), as calculated by the formula below, represents the ratio of the reduction in RT from the low-probability location (LPL) to the high-probability location (HPL) relative to the RT at the LPL: *P =* (RT_LPL_-RT_HPL_) / RT_LPL_. A larger value of *P* indicates a stronger learned suppression effect.

## 3 Results

### Distractor-absent trials

As shown in Figure 2A, the main effect of location was not significant for accuracy (*b* = 0.06, *SE* = 0.08, *z* = 0.72, *p* = 0.47, 95% CI from −0.102 to 0.220, *OR* = 1.06). However, the main effect of location was significant for response time (*b* = −19.10, *SE* = 2.84, *t*_(6655)_ = −6.71, *p* < 0.001, 95% CI from −24.67 to −13.52; *β* = −0.07, 95% CI from −0.10 to −0.05). Participants showed slower responses when the target appeared at high-probability distractor locations (785.09 ± 19.41 ms) than at low-probability distractor locations (746.36 ± 14.44 ms). These results suggest a robust learned spatial suppression effect at high-probability locations.

**Figure 2.**
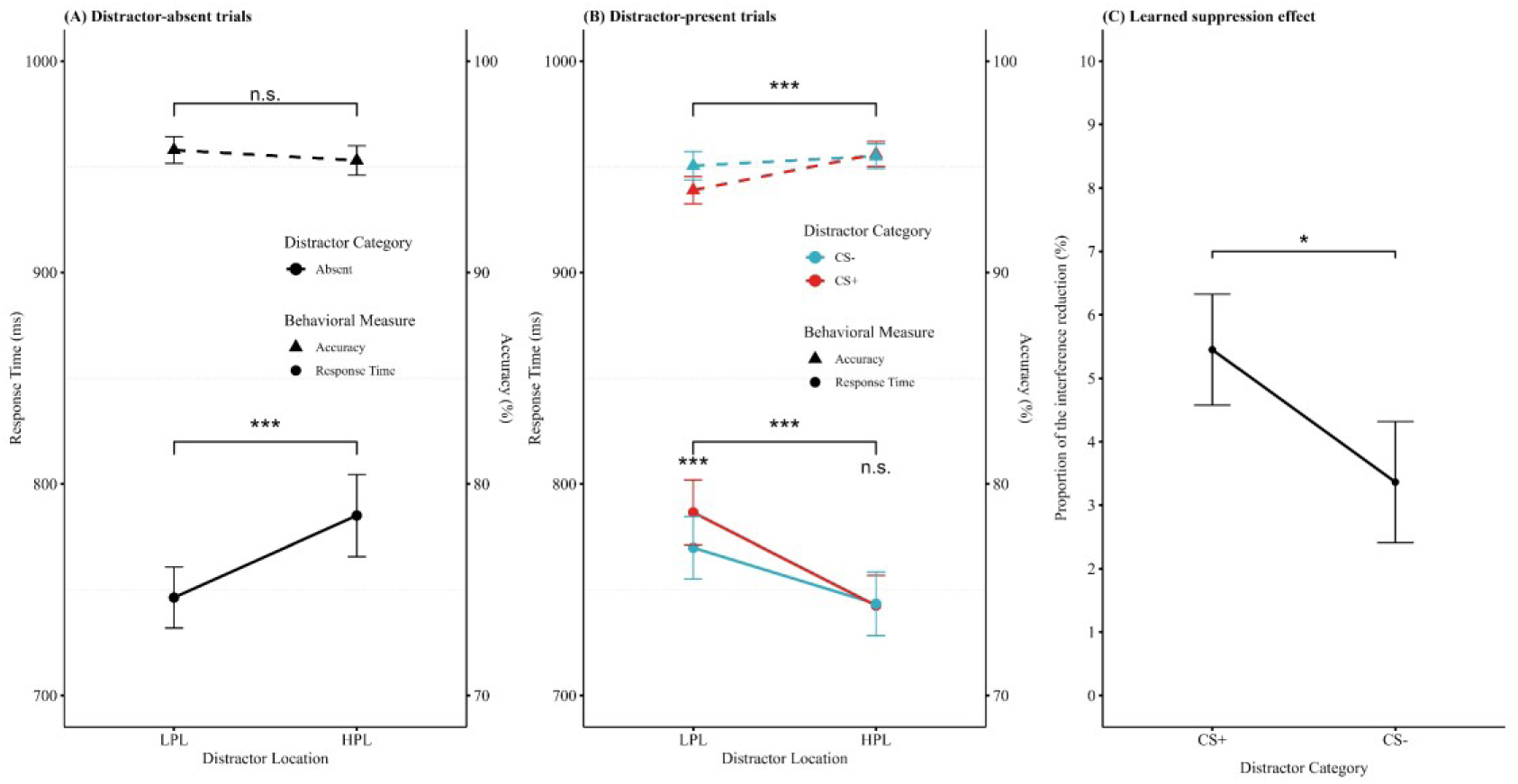
Learned spatial suppression is stronger when threat-history distractors appear at high-probability locations. *Note.* Behavioral results for visual search with distractor-absent trials (A) and distractor-present trials (B). The left Y-axis shows reaction time, represented by solid circles, and the right Y-axis shows accuracy, represented by solid triangles. (C) The proportion of reduction in interference (from the LPL to the HPL) for different distractor category. All error bars represent the within-subject standard error of the mean. LPL: low-probability distractor locations; HPL: high-probability distractor locations; CS+: threat-history distractors; CS-: no-threat-history distractors. *** indicates *p* < 0.001, * indicates *p* < 0.05, and n.s. indicates not significant (*p* > 0.05). See the online article for the color version of this figure.

### Distractor-present trials

For accuracy, as shown in Figure 2B, there was a significant main effect of distractor location probability (*b* = −0.11, *SE* = 0.04, *z* = −3.29, *p* < 0.001, 95% CI from −0.18 to −0.05, *OR* = 0.89). Higher accuracy was noted when the distractor appeared at high-probability locations compared to low-probability ones. However, the main effect of distractor category was not significant (*b* = −0.05, *SE* = 0.04, *z* = −1.48, *p* = 0.14, 95% CI from −0.12 to 0.02, *OR* = 0.95). The interaction between distractor location probability and distractor category was marginally significant (*b* = −0.06, *SE* = 0.04, *z* = −1.81, *p* = 0.07, 95% CI from −0.13 to 0.01, *OR* = 0.94). Follow-up analyses revealed that accuracy was significantly higher when threat-history distractors appeared at high-probability distractor locations than at low-probability distractor locations (HPL: 95.61 ± 0.59% vs. LPL: 93.90 ± 0.65%; *b* = −0.35, *SE* = 0.10, *z* = −3.71, *p* = 0.001).

For response time, the main effect of distractor location was significant (*b* = 17.68, *SE* = 1.33, *t*_(16606)_ = 13.33, *p* < 0.001, 95% CI from 15.08 to 20.28; *β* = 0.09, 95% CI from 0.08 to 0.11). Participants responded faster to the target when the distractor appeared at high-probability distractor locations than at low-probability distractor locations (*b* = 35.40, *SE* = 2.65, *z* = 13.33, *p <* 0.001). The main effect of distractor category was also significant (*b* = 3.92, *SE* = 1.33, *t*_(16606)_ = 2.95, *p* = 0.003, 95% CI from 1.32 to 6.52; *β* = 0.02, 95% CI from 0.01 to 0.04). Threat-history distractors elicited slower responses to the target than no-threat-history distractors (*b* = 7.84, *SE* = 2.65, *z* = 2.96, *p* = 0.003). As well as a significant interaction effect between distractor location and distractor category (*b* = 4.39, *SE* = 1.33, *t*_(16606)_ = 3.31, *p* < 0.001, 95% CI from 1.79 to 6.99; *β* = 0.02, 95% CI from 0.01 to 0.04). Further analysis revealed slower response to the target when threat-history distractors appeared at low-probability distractor locations compared to no-threat-history distractors (CS+: 786.53 ± 15.38 ms vs. CS−: 769.89 ± 14.75 ms; *b* = 16.61, *SE* = 4.29, *z* = 3.87, *p* < 0.001). However, no difference was observed at high-probability locations (CS+: 742.54 ± 14.24 ms vs. CS−: 743.38 ± 15.07 ms; *b* = −0.93, *SE* = 3.13, *z* = −0.30, *p* = 0.99). There was a difference in response to the target when distractors appeared at high-probability distractor locations compared to low-probability distractor locations (CS+: *b* = 44.14, *SE* = 3.76, *z* = 11.73, *p* < 0.001; CS−: *b* = 26.60, *SE* = 3.74, *z* = 7.11, *p* < 0.001).

### Learned suppression effect

As shown in Figure 2C, the results indicted that the proportion of interference reduction for threat-history distractors was significantly higher than that for no-threat-history distractors ( CS+: 5.45 ± 0.87% vs. CS−: 3.36 ± 0.95%; *t* _(29)_ = 2.19, *p* = 0.04, *d* = 0.40).

### Awareness of statistical regularities

Ten of the 30 participants reported not noticing whether the distractor appeared more or less frequently at a specific location. These participants reported an average confidence rating of 4.40 ± 0.50 on the 7-point scale. The remaining 23 participants reported noticing a spatial imbalance, with an average confidence rating of 4.26 ± 0.32. However, none of these participants correctly identified the high- or low-probability distractor locations as defined in the experiment. These results suggest that the learned suppression effect was driven by implicit awareness.

## 4 General Discussion

This study used threat conditioning task to create stimuli with or without threat history. In a subsequent visual search task, distractor location probability was manipulated to examine how threat history influences distractor suppression. The results show that faster target responses when distractors appeared at high-vs. low-probability locations, with stronger suppression for threat-history distractors.

First, this study essentially replicates Theeuwes et al. (2025) with moderate modifications to the experimental procedure. Consistent with previous research, We found that threatening distractors can be effectively suppressed even in the absence of a pre-conditioning phase. Participants responded faster to targets when a threat-history distractor appeared at high-probability location than at low-probability location. However, in contrast to Theeuwes et al. (2025), we observed a significant interaction between distractor category and location probability. One possible explanation is that Theeuwes et al. (2025) continued to administer shocks during the post-conditioning phase to maintain threat associations. The sustained emotional arousal may have disrupted the updating of suppression mechanisms, thereby attenuating the interaction between threat and location probability(Anderson & Britton, 2020). For example, Anderson and Britton (2020) found that, even though fixating on a threat-associated stimulus (CS+) would trigger an electric shock, participants still tended to direct their attention to this stimulus first. This result suggests that sustained emotional arousal may impair the adaptive attentional mechanisms, making it difficult for individuals to suppress the automatic attentional bias toward threat-related stimuli.

Furthermore, the present study provides additional evidence that adaptive attentional suppression mechanisms not only extend beyond physical salience (Failing & Theeuwes, 2020; Gong & Theeuwes, 2021), but also include motivational salience. Recent research has shown that salient distractors associated with fear, such as spiders or fear-conditioned stimuli, can be suppressed through statistical learning (Theeuwes et al., 2025; Theeuwes & van Moorselaar, 2025). Extending upon previous research, our study shows that threat-history distractors elicit stronger suppression effects than no-threat-history distractors. This difference emerges from the distinct influences of threat-related distractors at high- and low-probability locations. Specifically, our results support the idea that the high-probability location is covered by ‘blanket’ suppression, which prevents any item, regardless of its threat status, from capturing attention. In contrast, threatening stimuli are more likely to capture attention and impair visual search at other locations. Additionally, prior studies indicate that controllability modulates attentional bias toward threat stimuli (Bishop et al., 2004; Limbachia et al., 2021; Wood et al., 2015). When threat-related distractors repeatedly appear at predictable locations, increased controllability may reduce their attentional capture. These findings align with the notion that the visual system integrates statistical learning and motivational salience into a unified priority map (Gaspelin et al., 2025; Gaspelin & Luck, 2018; Pearson et al., 2022). Furthermore, our findings highlight the flexibility of attentional suppression, which is shaped by prior motivational experiences and reflects the combined influence of multiple adaptive attentional mechanisms(Anderson, 2021).

Taken together, the present findings provide evidence that threat-history distractors are more effectively suppressed when they appear in a predictable location via an adaptive attentional suppression mechanism. However, several important questions remain. Future studies could use neural measures, such as event-related potentials (e.g., P_D_ or N2pc) or eye-movement, to track the development of suppression over time and determine whether threat history affects early attentional selection or later-stage suppression processes(Gaspelin et al., 2023; Lin et al., 2024; Nobre & Ede, 2023; Zhang et al., 2025). Additionally, it would also be informative to explore whether passive exposure to threat-related stimuli, such as in habituation paradigms(Won & Geng, 2020), elicits similar modulation of learning-based suppression. Addressing these questions will help delineate the mechanisms through which threat history shapes attentional control.

## Ethics approval

The study was approved by the Committee on Human Research Protection of the School of Psychology, Guizhou Normal University (IRB approval number: GZNUPSY.NO2022E [007]).

## Consent to participate

All participants provided informed consent in accordance with the Declaration of Helsinki.

## Consent for publication

The article has been approved by all authors for publication.

## Availability of data and materials

All data supporting the findings are openly available via the Science Data Bank (https://www.scidb.cn/anonymous/VlpqNm55).

## Code availability

The analysis code supporting this study will be made publicly available on the Science Data Bank after publication.

## Authors’ contributions

Jingqing Nian served as lead for data curation, formal analysis, and validation and contributed equally to conceptualization, writing–original draft, and writing–review and editing. Yu Zhang served as lead for conceptualization, supervision, and writing– review and editing. Yu Luo served as lead for conceptualization, funding acquisition, supervision, writing–original draft, and writing–review and editing.

## Acknowledgements

This research was supported by the Basic Research Program of Guizhou Province (Qiankehe Jichu -ZK[2023] General-276) and the National Natural Science Foundation of China (32260209).

